# Tissue scarring provides a biomechanical framework to promote mammalian bile duct regeneration through the activation of integrin-SRC/FAK signalling

**DOI:** 10.1101/2025.04.21.649531

**Authors:** Alexander Walker, Paula Olaizola, Euan Brennan, Edward J Jarman, Yuelin Yao, Elizabeth Carmichael, Andreea Gradinaru, Alexander EP Loftus, David H Wilson, Anabel Martinez Lyons, Laura Charlton, Kimberley Ober-Vliegen, Wunan Mi, Amy Broeders, Kyle Davies, Neil O. Carragher, Asier Unciti-Broceta, Timothy J Kendall, Luc van der Laan, Monique MA Verstegen, Margaret C Frame, Scott H Waddell, Luke Boulter

## Abstract

Following chronic injury, the adult mammalian bile duct regenerates by forming new branches, essentially replumbing the ductular system to overcome blockages and breaks. To regenerate effectively, biliary epithelial cells (BECs) receive a range of pro-mitogenic signals from myofibroblasts, which concurrently deposit a collagen-rich scar around the duct as it regrows. Despite epithelial regeneration and scarring occurring side-by-side, whether the deposition of scar tissue regulates ductular regeneration *per se* remains unclear.

By inducing ductular fibrosis and regeneration *in vivo*, we show that the formation of collagen-I-rich scars around regenerating ducts changes the local biomechanical properties of these tissues, promoting the growth of ducts. Critically, this changing structural landscape is perceived by a spatially restricted population of biliary epithelial cells which forms a “leading-tip” of integrin-α2-high cells. This leading-tip undergoes partial-EMT-type reprogramming, allowing it to become migratory and coordinate ductular regeneration. We show that this process is directly driven through an integrin-α2-SRC/FAK signalling axis; thereby connecting epithelial regeneration directly to the changing fibrotic environment in chronic ductular disease.

**Highlights:**

- Chronic liver disease results in the formation of stiff, collagen scars around ducts.
- New ducts acquire high levels of integrin-α2 which is spatially localised to a “leading-tip”, which loses epithelial features.
- Integrin-α2β1-SRC/FAK signalling regulates ductular migration by linking changes in the bio-structural composition of the liver with ductular cells.

## Introduction

During chronic liver disease, the development of chronic fibrosis and liver scarring is synonymous with tissue regeneration, where normally quiescent, terminally differentiated epithelial cells re-enter cell cycle^1–5^. While this concept has long been accepted, how these two tissue level processes (fibrosis and epithelial proliferation/regrowth) interact to restore a correctly pattered tissue following injury is less clear.

In ductular liver diseases, such as primary sclerosing cholangitis, where bile ducts become broken, strictured or completely blocked, biliary epithelial cells (BECs) proliferate to form collateral ducts which re-constitute the biliary network and restore tissue function, thereby resuming bile flow^6^. This is not a trivial process, requiring not only the production of new BECs through proliferation, but also the collective migration of ductular cells as a tubular structure, with correct spatial and geometric orientations to form a new duct branch within a solid organ^7^. These ducts are surrounded by myofibroblasts, which provide pro-regenerative signals including JAGGED-1 and Hedgehog ligands, which drive BEC proliferation^8–10^. Concurrently, duct-associated myofibroblasts produce a collagen-rich extracellular matrix which evolves with ductular regrowth^11^. During tissue embryogenesis and adult tissue morphogenesis (particularly in the mammary duct), the correct deposition of collagen controls how ductular systems develop^12,13^. In these growing systems, the rigidity, complexity, and alignment of collagen fibrils instructs groups of ductular cells to establish where a new ductular branch will form and dictates the ultimate dimensions of the duct that will grow. Despite the close association of collagen and BECs in ductular regeneration in the liver, whether the deposition of collagen during ductular scarring regulates how new bile ducts form during regeneration has not been explored.

During tumour initiation, stiffer extracellular microenvironments promote cancer cell proliferation and metastasis through activation of mechanosensitive signalling pathways, including several kinases (e.g., SRC and FAK)^14^. The major receptor family that perceives changes in extracellular matrices are integrins, plasma-membrane bound heterodimeric receptors, which have variable substrate specificity determined by which receptor sub-units are present^15^. As such, integrins interpret changes in matrix macromolecules, such as collagens, laminins and fibronectin, as well as matrix-bound growth factors like latent-TGFβ, which becomes activated upon interaction with αV-containing integrins^16^.

Work in polycystic liver disease, where bile ducts form pathological cysts, and in bile duct cancer (cholangiocarcinoma) has shown that integrin-sensing of the extracellular matrix is a critical factor in disease progression^17,18^. We hypothesised, therefore, that during ductular regeneration BECs sense and respond to their changing physical landscape through integrins and that this extracellular matrix (ECM)-BEC signalling is essential for ductular regeneration. Here, we show that during bile duct disease the biophysical nature of the liver changes as it scars. Ductular cells respond to this new environment by upregulating specific integrin receptors within anatomically constrained populations of BECs, forming a leading-edge (analogous to tip-cells during sprouting angiogenesis). Integrin-signalling activates a range of downstream kinases including SRC, FAK and EPHA2, which drives a partial-EMT enabling BECs to migrate through tissues, thereby driving ductular regeneration.

## Results

### Dynamic remodelling of the bile duct extracellular microenvironment is a hallmark of ductular regeneration

The activation of fibroblasts and deposition of scar-forming collagens is a hallmark of chronic disease in a variety of paediatric and adult bile duct pathologies, where scars surround and form close spatial associations with regenerating BECs^1,19^. While it is known that fibroblast-BEC interactions are important for ductular regeneration, many of the regulatory, pro-regenerative signals which mediate adult bile duct repair have been identified due to their role in ductular development^20^. Jagged-Notch signalling (an archetypal signalling pathway in ductular ontogeny), for example, is essential for adult bile duct regeneration, where portal fibroblasts produce Jagged-1 and activate Notch1/2 on regenerating BECs^2,8,21,22^.

Using a published single cell data set from normal human liver or from patients with chronic ductular disease^23^ (primary sclerosing cholangitis, PSC; and primary biliary cholangitis, PBC) we sought to identify *de novo*, disease-specific interactions that occur between fibroblasts and BECs.

Following identification of all the major liver cell populations using SingleR which performs unbiased cell type recognition from single cell data^24^ (**Supplementary Figure 1A**) we validated the presence of fibroblasts (defined by the expression of *FAP*, *ACTA2* and *PDGFRA*) and BECs (through the presence of *EPCAM*, *SPP1* and *SOX9* transcripts) within these data (**Supplementary Figure 1B and 1C**). Using CellChat ligand-receptor interaction modelling^25^, we identified 55 putative receptor-ligand interactions between fibroblasts and BECs in normal, healthy tissue **(Figure 1A** and **Supplementary Table 1**). In PSC/PBC patients the number of predicted interactions between these same cell types increased to 308, suggesting that in biliary disease the global signalling repertoire of both fibroblasts and BECs is increased. As expected, CellChat identified JAGGED1-NOTCH2 and JAGGED1-NOTCH3 interactions between fibroblasts and BECs. Additionally, we found a large number of candidate interactions between *COL1A1* expressed by the fibroblast population and GFOGER-binding integrins which are considered receptors for a range of ECM proteins, including collagen-I (**Supplementary Table 1**). To understand where collagen-I-binding integrins are expressed, we collated a GFOGER-integrin gene signature looking at the combined expression of *ITGA1*, *ITGA2*, *ITGA10*, *ITGA11* and *ITGB1*, and when we show this signature in single cell data from healthy and PSC/PBC liver, BECs, Fibroblasts and Endothelial cells are all enriched for the expression of integrins, which are known to interact with collagens (**Figure 1B**). Collectively, these data indicate that the presence of ECM proteins within PSC and PBC have the potential to regulate how adult ducts regenerate.

**Figure 1:**
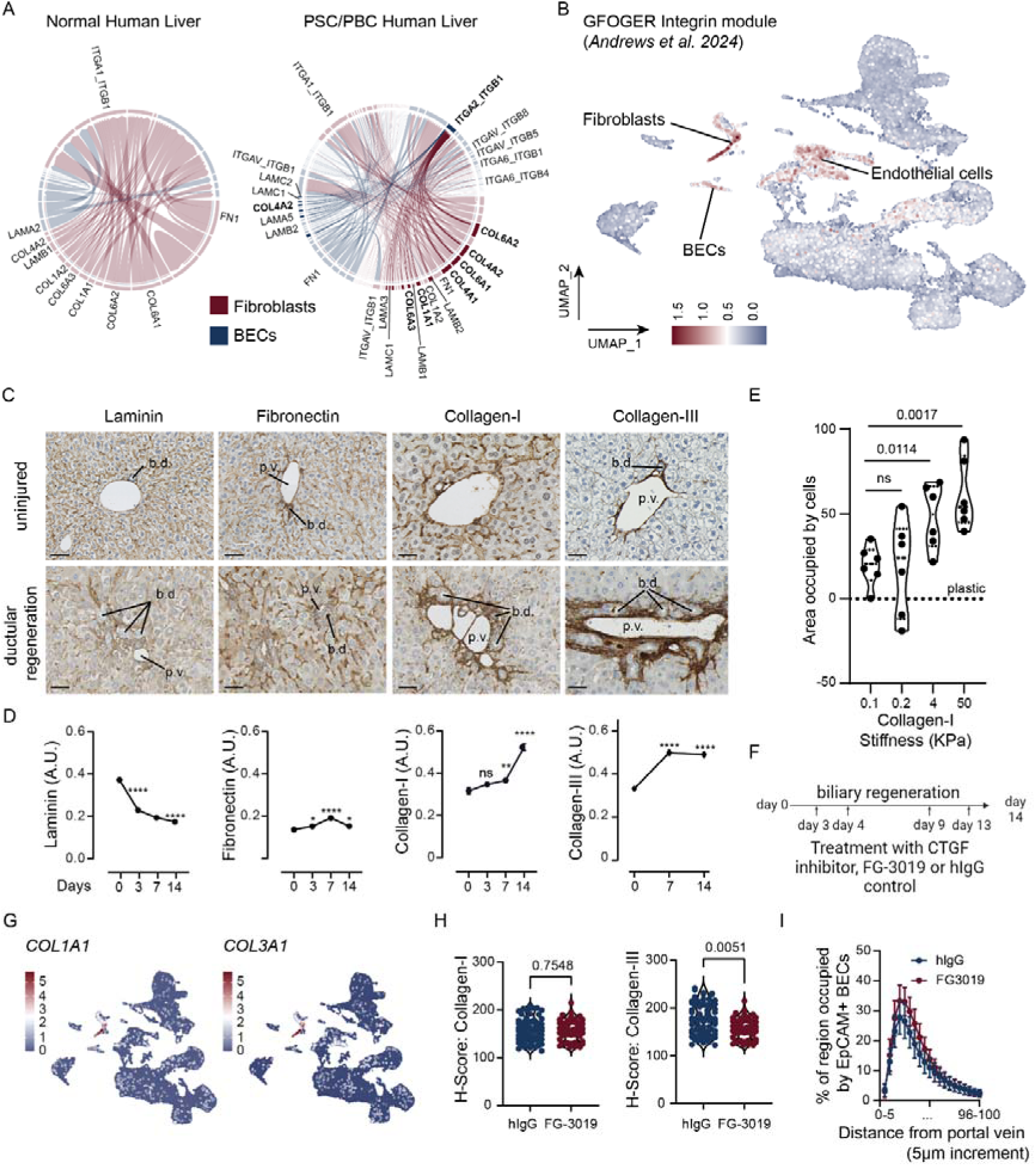
Cholestatic disease is typified by the formation of new ECM-BEC interactions. **A.** CellChat ligand-receptor interaction analysis between fibroblasts (blue) and BECs (red) in normal human liver and PSC/PBC human liver. **B.** UMAP of single cell RNA sequencing of normal liver or liver from patients with primary sclerosing cholangitis (PSC) or primary biliary cholangitis (PBC) highlighting Fibroblasts, Endothelial cells, and Biliary Epithelial Cells (BECs) showing the combined expression of GFOGER-binding integrins. **C.** Immunohistochemistry of laminin, fibronectin, collagen-I and collagen-3 from uninjured mouse liver (top panels) and following 14 days of DDC-induced ductular regeneration (lower panels) scale bar = 100µm. b.d. bile duct, p.v. portal vein. **D.** Quantification of ECM immunohistochemistry over progressive DDC-induced ductular regeneration (N=4 individual mice with n=10 lobules quantified per animal). **E.** Cell growth of human BECs on collagen-I of increasing stiffnesses (from 0.1 to 50KPa) n=6 technical replicates. Data is normalised to culture on tissue culture plastic, dotted line. **F.** Schematic of inhibiting CTGF during DDC-induced bile duct regeneration using FG-3019 or hIgG control. **G.** COL1A1 and COL3A1 mRNA expression in human normal, PSC/PBC single cell data**. H.** Quantification (H-score) of peri-portal collagen-I and collagen-III following treatment with FG-3019 or hIgG (N=5 mice with n=10 regions samples per animal)**. I.** Distance of migrating EpCAM-positive ducts from the portal vein following FG-3019 (N=7 individual animals) or hIgG (N=8 individual animals) treatment during ductular regeneration.

Patients with PSC and PBC experience progressive fibrosing disease throughout their lives^26^. To model the progressive damage and stricturing of bile ducts and concomitant regeneration of the biliary tree, we administered 3,5-diethoxycarbonyl-1,4-dihydrocollidine (DDC) to mice resulting in the formation of porphyrin-rich plugs within the duct lumen, blocking normal bile flow^27^. Following the initiation of injury, the levels of laminin, a key component of the basement membrane surrounding the uninjured bile duct, are rapidly reduced (within the first 3 days of ductular regeneration), while the levels of periductular fibronectin remain relatively static. Analogous to human disease, we also found that the amount of collagen-I and collagen-III proteins deposited around regenerating bile ducts increases as disease progresses suggesting that the transition from a laminin-to-collagen rich environment may promote ductular regeneration (**Figure 1C** and **1D**).

While both collagen-I and -III are found adjacent to regenerating ducts, it is unclear whether they share function or whether one ECM component is structurally dominant over the other. Indeed, in other tissues wound healing relies on the early deposition of collagen-III which is more elastic only to be replaced by a more tensile network of collagen-I^28^. To test whether collagen stiffness dictates BEC proliferation, the human cholangiocyte cell line, H69 BECs (hereafter known as hBECs), were plated on collagen-I of different physiological (0.1-4KPa) and supraphysiological (50KPa) stiffnesses. Across stiffnesses, hBEC proliferation increases compared to cells which are not plated on collagen-I. Furthermore, as stiffness of collagen-I increases, so too does the proliferation of hBECs suggesting that both the presence and stiffness of collagen-I influences their growth (**Figure 1E**). Connective tissue growth factor (CTGF) secreted by hBECs promotes adjacent fibroblasts to produce collagen^11^. We initiated ductular regeneration during which animals were treated *in vivo* with a CTGF-neutralising antibody^29^, FG-3019 or hIgG isotype control (**Figure 1F**). In addition to COL1A1, COL1A3 is highly expressed by PSC/PBC fibroblasts, however CellChat failed to identify any interactions driven by the presence of COL3A1 (**Figure 1G**). Inhibition of CTGF did not affect the levels of peri-ductular collagen-I, however significantly reduced the levels of peri-ductular collagen-III (**Figure 1H**).

During duct regeneration, nascent ductules form from parental ducts in a process analogous to blood vessel collateralisation, where new ducts migrate into the surrounding tissue to restore tissue function. To assess ductular migration *in vivo,* we utilised the portal vein as an anchor and assessed how far ducts were found away from these structures (**Supplementary Figure 1D and 1E**), failure to migrate would lead to an accumulation of ducts close to the vein and enhanced migration would result in ducts further from the vein; using this method we are able to accurately demonstrate that as ductular regeneration progresses (between day 0/uninjured and day 14) that Keratin-19 positive ductules are found to progressively migrate into the liver tissue (**Supplementary Figure 1F**). We measured the migration of regenerating ductules in mice treated with CTGF or hIgG and found that the migration of ducts into the tissues does not change following CTGF-inhibition (**Figure 1I**), suggesting that collagen-III levels *in vivo* are dispensable for normal ductular migration.

### Integrin-α2 demarcates emerging ducts during ductular regeneration

Integrin receptors exist as αβ-heterodimers which sense changes in both the biochemical and mechanical composition of the extracellular matrix. In order to alter their substrate specificity and perceive changes in extracellular matrix proteins, β1-integrins bind to a diverse range of α-integrin subunits. In human PSC/PBC, ligand-receptor interaction prediction identified COL1A1-ITGA2-ITGB1 interactions as specific to PSC/PBC (**Figure 1A** and **Supplementary Table 1**) and while BECs from PSC patients express the same level of *ITGB1* mRNA as those from normal liver, the transcriptional expression of *ITGA2* and protein level of ITGA2 in PSC BECs is increased (**Figure 2A** and **2B, Supplementary Table 2**).

**Figure 2:**
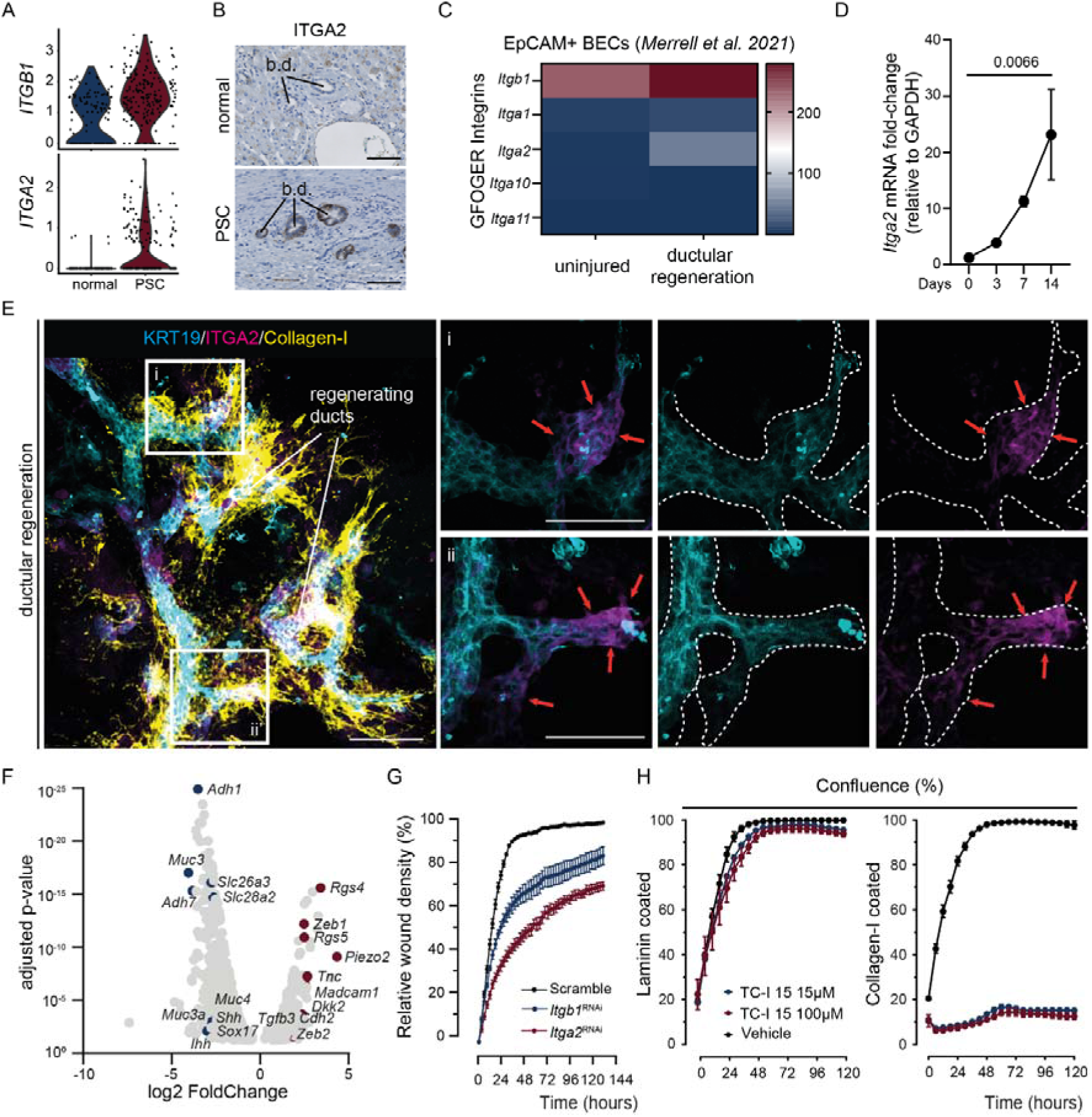
Integrin-α2 defines a migratory, pro-regenerative population of BECs. **A.** Violin plots demonstrating the transcriptional expression of *ITGB1* and *ITGA2* in BECs from scRNA sequencing of normal liver (blue) and from patients with Primary Sclerosing Cholangitis (burgundy), derived from Andrews et al 2024. **B.** Immunohistochemistry of ITGA2 in normal human liver and liver from Primary Sclerosing Cholangitis. Scale bar = 100µm. **C.** mRNA expression of GFOGER-binding integrins in EpCAM+ BECs 28 days following DDC-treatment, derived from Merrell et al. 2021. **D.** mRNA of Itga2 mRNA expression over the time course of DDC injury. **E.** FUnGI imaging of regenerating bile ducts showing KRT19+ BECs (cyan) ITGA2 (magenta) and collagen-I (yellow). Scale bar = 200µm. **F.** Volcano plot of differentially expressed genes from ITGA2-high versus ITGA2-low lineage traced BECs (N=3 mice). **G.** Migration of hBECs treated with a non-targeting (scrambled) RNAi or RNAi specifically targeting *Itgb1* or *Itga2*. N=3 experimental replicates. **H.** Confluence of hBEC when cultured on either laminin (left) or collagen-I (right) in the presence of 15µM or 100µM TCI-15 (N=3 experimental replicates). b.d.: bile duct.

The utilisation of integrin-α2 during ductular regeneration is conserved and during experimentally-induced ductular regeneration, bulk RNA sequencing analysis of EpCAM-positive cholangiocytes^30^ demonstrated increased expression of the GFOGER-binding integrins *Itgb1* and partners *Itga1* and *Itga2* (but not *Itga10* and *Itga11,* **Figure 2C**). The mRNA expression of *Itga2* increases over the time course of ductular injury and regeneration (**Figure 2D**), whereas the expression of other ECM-binding (*Itga1* and *Itgb1*) and TGFβ-activating (*Itgav*, *Itgb6* and *Itgb8*) are more dynamic as regeneration progresses (**Supplementary Figure 2A**). As we have previously reported, we failed to find any expression of ITGA2 in the healthy mouse bile duct (whilst ITGB1 is constitutively expressed)^17^; however, within 7 days of DDC-induced regeneration small ductules and isolated ductular cells are positive for ITGA2 protein and by 14 days large numbers of regenerating BECs express ITGA2 (**Supplementary Figure 2B**), and while ITGA1 is widely expressed by sinusoidal endothelial cells, it is largely absent in BECs (**Supplementary Figure 2C**). While BECs upregulate *Itga2* expression, in patients with PSC the protein levels of ITGA2 vary both between and within patient samples (**Supplementary Figure 2D**) suggesting that regionalisation of integrin signalling could be important in coordinating ductular regeneration.

To further explore the spatial heterogeneity of integrin-α2 in the biliary tree we used FUnGI tissue clearing to image regenerating bile ducts^17,31^. Rather than being widely expressed throughout the duct, 14 days following the start of regeneration, ITGA2 is enriched in newly forming “leading-tip” cells which branch-off mature Keratin-19 positive ducts and which are surrounded by an expansive network of collagen-I (**Figure 2E**).

There is limited evidence for a *bona fide* stem cell in the regenerating adult liver; rather, BECs undergo epigenetic remodelling, enabling them to enter a pro-regenerative cellular state^4,6^. We propose that ITGA2-positive, regenerating ducts necessarily differ from parental ducts, acquiring features that enable them to lost their normally polarised phenotype and migrate as a duct to recover ductular function. To understand this further we irreversibly labelled BECs *in vivo* with the fluorophore tdTomato (by treating *Krt19CreER^T^; R26^LSL-tdTomato^* transgenic mice with tamoxifen) and then initiating ductular regeneration. We isolated EpCAM+/tdTomato+/ITGA2+ BECs (which had been labelled and then transitioned into a “regenerative” ductular state) or EpCAM+/tdTomato+/ITGA2-BECs (those which had been labelled and not entered a regenerative state) from DDC-injured livers. Lineage-traced, ITGA2-positive BECs are transcriptionally enriched for genes associated with EMT and migratory-like phenotypes in other tissues (**Supplementary Table 3**), including the EMT transcriptional regulator *Zeb1, Cdh2* (which encodes for N-cadherin) and *Tnc* (a matricellular protein known to support EMT) (**Figure 2F**).

Our data suggest that the formation of a collagen-I rich extracellular matrix regulates the migratory phenotype of BECs. Indeed, hBECs treated with RNAi against either *Itga2* or its obligate binding partner *Itgb1* results in significantly reduced migration *in vitro* (**Figure 2G**). Critically, our data suggests that ECM substrate is essential for the acquisition of this migratory phenotype and when hBECs are treated with a selective inhibitor of the integrin-α2β1 heterodimer, TC-I 15^32^, cellular migration is only inhibited when cells are grown on collagen-I and not when they are grown on laminin (**Figure 2H and Supplementary Figure 3A-C**).

### Integrin-α2β1 alters SRC:FAK activity in migrating BECs

Engagement of integrin heterodimers with ECM results in the formation of a multiprotein adhesion complex which triggers molecular signalling events through the activation of a range of intracellular intermediates. To rationalise which signalling pathways are activated downstream of integrin-α2β1, we silenced the expression of either *Itga2 or Itgb1* using RNAi in hBECs (or pharmacologically disrupting integrin-α2β1 function by treating hBECs with TC-I 15) and using multiplex reverse phase protein arrays to simultaneously assay 80 (phospho)proteins identified that the ratio of pFAK^Y397^:FAK and pAKT^S473^:AKT were significantly supressed (compared to scrambled RNAi sequences) when *Itga2* and *Itgb1* are lost (**Figure 3A, Supplementary Figure 3D**).

**Figure 3:**
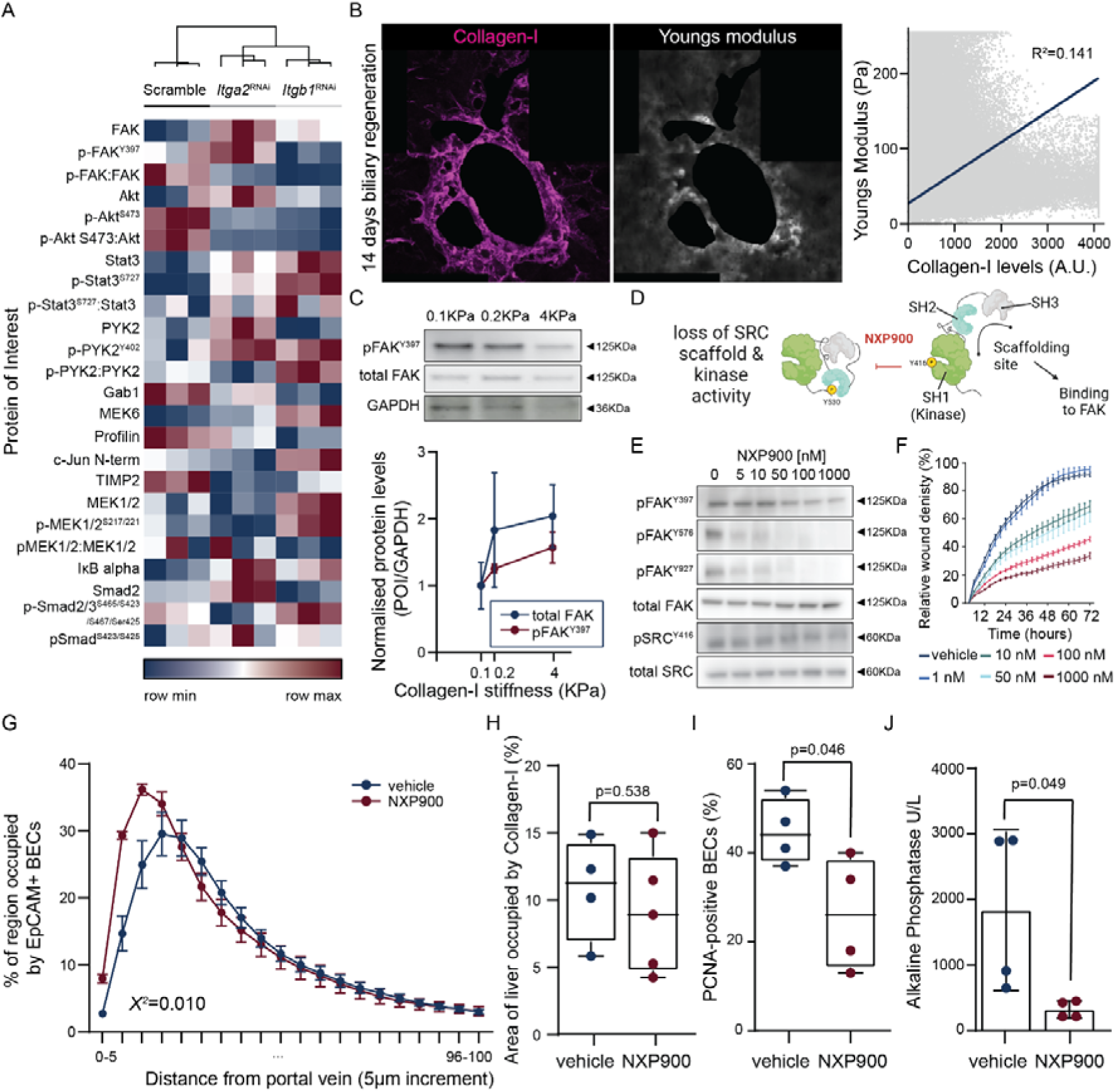
Integrin-α2β1 mediates migration through SRC:FAK activation. **A.** Reverse Phase Protein Array demonstrating changes in (phospho)protein levels in hBECs treated with non-targeting scrambled, *Itga2* or *Itgb1* RNAi (N=3 experimental replicates). **B.** Confocal imaging of collagen-I (magenta, left) and AFM (right). Correlation plot between collagen-I intensity and Youngs modulus (stiffness) **C.** Immunoblotting (upper panels) and quantification (lower panel) showing the changes in total FAK and phosphorylated FAK^Y397^ from BECs plated on different stiffnesses of collagen-I. **D.** Schematic demonstrating the mode of action of NXP900. **E.** Immunoblotting demonstrating the levels of total or phosphorylated FAK (FAK^Y397^, FAK^Y576^, FAK^Y927^) and total or phosphorylated SRC (SRC^Y416^) with increasing concentrations of NXP900. **F.** Migration of hBECs following SRC:FAK inhibition with NXP900 (n=4 technical replicates per treatment). **G.** Migration of regenerating ducts following treatment with NXP900 or vehicle *in vivo*. (N=5 individual animals per group) **H.** Quantification of collagen-I positive area (N=4-5 individual animals per group) and **I.** PCNA/EpCAM-positive BECs following NXP900 treatment (N=4 animal per group)**. J.** Serum Alkaline Phosphatase and Bilirubin levels following NXP900 treatment (N=4 individual animals).

In liver cancer, the rigidity of ECM cell substrates promotes cancer cell proliferation^33^. Using atomic force measurements of the regenerating liver, we found that regionalised stiffness around regenerating bile ducts correlates with the expression of collagen-I (**Figure 3B**), implying that as new ducts emerge from the nascent biliary tree, they experience an altered biomechanical landscape. Indeed, when hBECs were plated on collagen-I of stiffnesses ranging from 0.1-4KPa the levels of total and phosphorylated FAK increases (**Figure 3C**). FAK activates both SRC (which in turn reciprocally phosphorylates FAK) and AKT to regulate cellular proliferation and migration. In hBECs, treatment with Pictilisib, a potent inhibitor of PI3K reduced the expression of pAKT^S473^, however failed to prevent the migration of hBECs *in vitro* or ductular migration *in vivo* (**Supplementary Figure 3E-G**). Conversely, using an inhibitor of SRC-FAK, NXP900, which prevents both the catalytic and scaffolding activities of SRC^34^, significantly reduced FAK activation (**Figure 3D and 3E**) and the migration of hBECs *in vitro* in a dose-dependent manner. To test whether SRC-FAK activity drives ductular regeneration *in vivo*, we initiated ductular regeneration and administered either NXP900 or vehicle alone for the duration of repair. NXP900-treated mice showed a robust inhibition of SRC phosphorylation at Y416 *in vivo* and concurrent loss of SRC-mediated phosphorylation of FAK at Y576/577, though autophosphorylation of FAK at Y397 is unaffected (**Supplementary Figure 3H**), as well as a significantly reduced ductular migration compared to vehicle-treated animals (**Figure 3G**), without affecting the levels collagen-I (**Figure 3H**). In addition to reduced ductular migration, NXP900 inhibition of SRC-FAK reduced the proliferation rate of BECs (**Figure 3I**) and levels of serum alkaline phosphatase (**Figure 3J**), demonstrating that SRC-FAK activation is required for normal ductular regeneration.

#### Integrin-α2β1 is necessary for BECs to regenerate mammalian bile ducts

The spatial distribution and activation of integrin-α2 during ductular regeneration suggests that signalling through this receptor plays a role in the formation of new ducts from the nascent, pre-existing biliary tree. Indeed, in mammary ducts, integrin-α2β1 is critical for normal ductular remodelling during pregnancy^35^, suggesting that facultative regeneration share common features across tissues.

BECs can be isolated from the adult liver and cultured as genetically stable intrahepatic cholangiocyte organoids (ICOs)^36^. ICOs are relatively simple hollow spheres in which BECs become transcriptionally homogenous throughout the time course of culture. When differentiated with R-spondin, Dexamethasone, EGF and DKK1, ICOs undergo morphological rearrangement, forming buds which elongate into ductular branches (**Supplementary Figure 4A and 4B**). These Branched Cholangiocyte Organoids (BRCOs) more closely reflect the complex tubular network seen in the mammalian liver. Using available single cell RNA sequencing data from ICOs and BRCOs, cells clustered into seven distinct Seurat clusters (**Figure 4A, Supplementary Figure 4C**), in which cluster 2 demonstrated the highest level of *ITGA2* expression (**Figure 4B**). Gene Set Enrichment Analysis of these seven Seurat clusters identified differentially enriched biological processes across groups of cells (**Supplementary Table 4**) with cluster 2 being enriched for a number of mitogenic and migratory WikiPathway terms (**Figure 4C** and **Supplementary Figure 4D**). By using the terms present in these WikiPathways that are expressed in BRCOs we generated aggregated modules of gene expression for Focal Adhesion (WP306), Integrin Mediated Cell Adhesion (WP185) and Hippo Merlin Signalling Dysregulation (WP4541); all of which are enriched within cluster 2 (**Figure 4D**), critically these signatures are all driven in part by the high levels of *Itga2* transcript expression (**Supplementary Figure 4E**).

**Figure 4:**
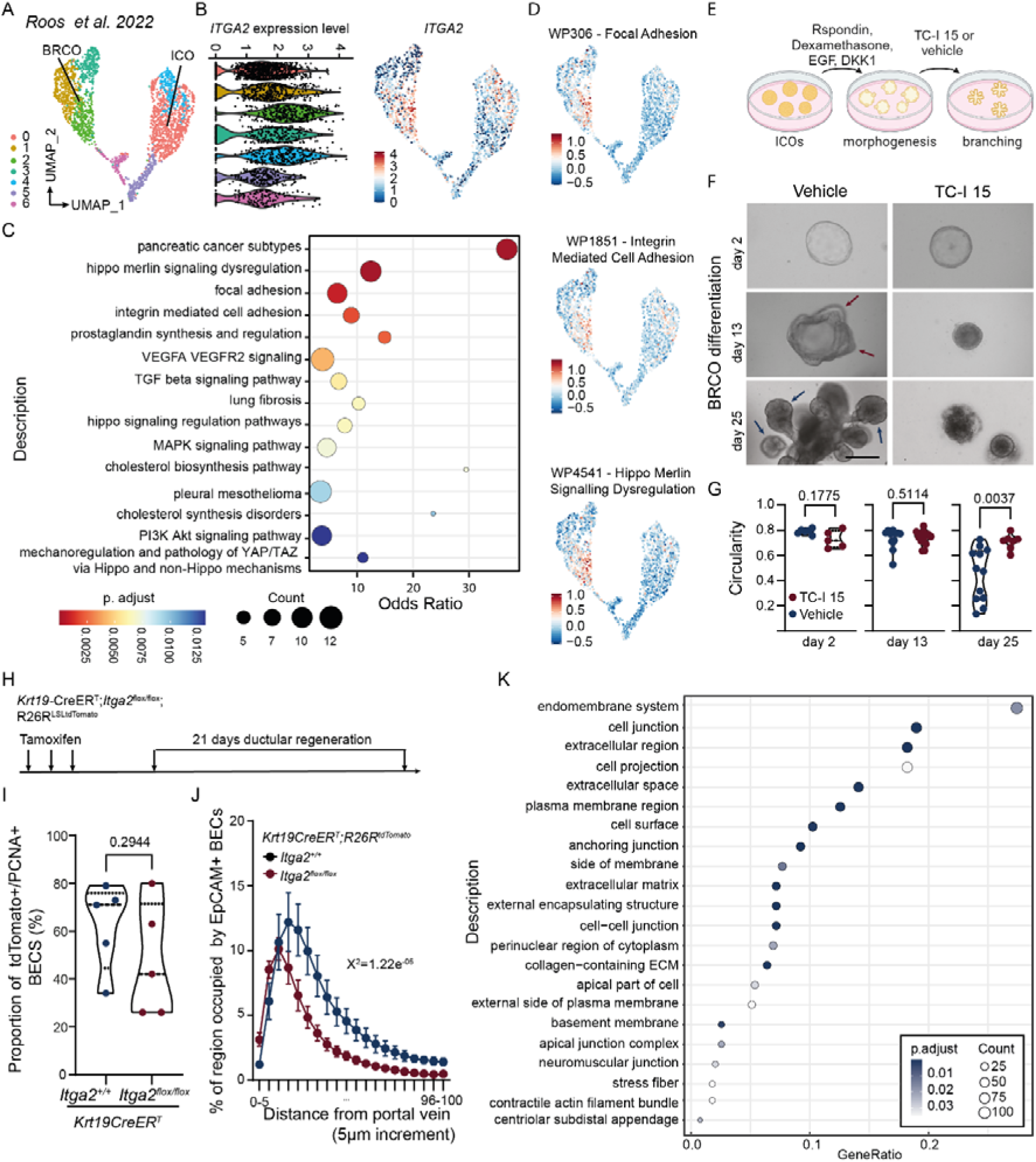
Integrin-α2β1 is essential for normal ductular regeneration. **A.** Seurat clusters of ICOs and BRCO scRNA sequencing *(Roos et al. 2022*). **B.** *ITGA2* mRNA levels in single cells across the seven Seurat clusters (left panel) and represented as a UMAP (right panel). **C.** Enrichment of WikiPathway terms within Cluster 2. **D.** UMAPs showing enrichment for WikiPathway terms Focal Adhesion, Integrin Mediated Cell Adhesion and Hippo Merlin Signalling Dysregulation within cluster 2. **E.** Schematic of the differentiation of Intrahepatic Cholangiocyte Organoids (ICOs) into Branching Cholangiocyte Organoids (BRCOs) in the presence of the integrin-α2β1 inhibitor, TC-I15. **F.** Photomicrographs of ICOs switched into BRCO differentiation media in the presence of TC-I 15 or vehicle alone at 2-, 13- and 25-days following differentiation. Red arrows denote thickening of the ICO wall and the formation of budding-like structures. Blue arrows identify the formation of established ductular branches (scale bar = 250μm). **G.** Histograms showing measurements of circularity, at 2-, 13- and 25-days of BRCO differentiation when treated with vehicle or TC-I 15. **H.** Schematic showing tamoxifen-induction of lineage tracing of *Itga2*^-/-^ BECs during DDC-induced ductular regeneration. **I.** Number of tdTomato-positive, lineage traced or PCNA+ tdTomato+ BECs in *Krt19CreER^T^*;*R26R^LSL-tdTomato^*;*Itga2*^flox/flox^ or *Itga2*^+/+^ mice (N=5 per group). **J.** Quantification of distance migrated by tdTomato-positive, lineage traced cholangiocytes with wild type *Itga2* or in which *Itga2* has been deleted (N=5). **K.** GOTerm analysis of bulk RNA sequencing from *Itga2^+/+^* or *Itga2^flox/flox^*tdTomato-positive cholangiocytes (N=4 per group).

Integrin-α2β1 is required for ducts to become cysts in polycystic liver disease^17^ and while the development of a collagen-I rich extracellular matrix is well described during ductular regeneration, whether GFOGER-sensing integrins play a functional role in perceiving this changing bio-mechanical landscape remains unclear. To explore whether BECs sense their collagen-I rich microenvironment we differentiated human ICOs into BRCOs in the presence of the selective integrin-α2β1 inhibitor, TC-I 15 (**Figure 4E**). Within the first two weeks of BRCO differentiation, organoids undergo morphogenic rearrangement where the organoid wall invaginates and thickens to form budding-like structures, reminiscent of early ductulogenesis. Over time these ductular structures become more complex and highly branched (**Figure 4F**, left panels). Following treatment with TC-I 15, however, ICOs remain spherical and do not form branches (**Figure 4F**, right panels). Furthermore, by measuring the branching levels of these ICOs we found that vehicle-treated organoids show changes in the number of branches that form at 13- and 25-days of differentiation, while TC-I 15-treated organoids show no substantial change to their branching during the timeframe of BRCO differentiation (**Figure 4G**).

Our data implicates integrin-α2β1 in the *de novo* formation of ducts; however, BRCOs are cultured in Matrigel, which does not have high levels of collagen-I observed *in vivo* and similarly lacks the instructive pro-mitogenic signals which are provided by activated portal fibroblasts and immune cells during ductular regeneration. To overcome this, we generated transgenic mice in which exon-1 of *Itga2* is flanked by loxP sites (*Itga2*^flox/flox^) and can be deleted in BECs by crossing these animals with *Krt19-CreER^T^;R26R^LSL-tdTomato^* mice^17,37^. *Krt19*-*CreER^T^;R26R^LSL-tdTomato^; Itga2*^flox/flox^ animals (or *Itga2*^+/+^ control animals) were dosed with tamoxifen to delete *Itga2* specially from BECs which show robust loss of the ITGA2 protein in tdTomato-positive lineage traced cells (**Supplementary Figure 4F**). Following bile duct regeneration, the number of lineage traced tdTomato-positive BECs in *Itga2*^+/+^ or *Itga2*^flox/flox^ mice remained the same and they demonstrated no differences in rate of proliferation (as determined by PCNA-positivity), suggesting that ITGA2-signalling is not mitogenic in BECs (**Figure 4I**). ECM-integrin signalling is known to mediate cellular migration and invasion across a number of cell and tissue types. Given new bile ducts undergo tubular migration in order to regenerate a functional ductular network, we measured the ability of tdTomato-positive BECs either with or without *Itga2* to migrate away from the portal tract *in vivo*. The median distance travelled by *Itga2*^+/+^ BECs was significantly higher than those lacking *Itga2,* (*Itga2^+/+^*: 16.03µm ± 9.39 vs *Itga2*^flox/flox^: 7.96µm ± 2.57) demonstrating that integrin-α2 is critical for BEC migration *in vivo*. Finally, *Itga2*^+/+^ or *Itga2*^flox/flox^ regenerating BECs were isolated by FACS and subject to bulk RNA sequencing. Analysis of differentially expressed transcripts showed enrichment for a range of enrichGO terms (**Figure 4K**) including those associated with cell junction, extracellular space, anchoring junction and cell-cell junction (**Supplementary Table 5**). Collectively, these data demonstrate that integrin signalling is required for regeneration of the mammalian biliary tree by perceiving the changing extracellular landscape in chronic biliary disease.

## Discussion

Chronic liver damage leads to two outcomes: regeneration to reconstitute the cellular mass of the organ or fibrosis which maintains the structural integrity of the tissue. Across the regenerating liver, these two processes co-exist; however, how the deposited extracellular matrices (scarring) interact with regenerating cells to co-ordinate faithful repair is poorly defined. Indeed, in tissues which are repopulated by resident stem or progenitor cells, such as the skin or intestine, the role of the extracellular matrix is critical for normal homeostatic maintenance and for injury-induced repair. During injury-induced, facultative regeneration in the liver, the composition of the extracellular matrix is dynamic and we show that a laminin-rich basement membrane is supplemented with fibrillar collagens-I and -III. The production of this extracellular matrix is conserved across species and these new structural components are almost exclusively produced by fibrogenic cells within the damaged liver. The deposition, cyclical reabsorption and re-deposition of hepatic scars is not a new concept; however, in the context of parenchymal regeneration, these processes have been seen as detrimental to regeneration, contributing to the formation of nodules in end-stage disease. Here we show that scarring, at least in the initiating phases of disease, is instructive providing both an altered biochemical and biophysical substrate which ductular epithelial cells utilise to exit the nascent duct forming a new collateral ductule in order to restore biliary function. Critically, we identify that these new ductules are characterised by the regionalised expression of integrin-α2, ensuring that they and not the parental duct can selectively sense the changes in collagen environment. Integrin-collagen interactions have been identified in several epithelial branching systems, particularly in the breast (during pregnancy), pancreas and lung (during development); these data demonstrate that the quantity, rigidity, and orientation of collagen fibres all contribute to the correct division and branching of mammalian ductular networks. Our data demonstrates that these physiological processes, which are critical for normal tissue patterning, are redeployed during ductular regeneration and that when considering how to develop anti-fibrotic strategies in chronic liver disease we should consider the context of cellular regeneration as these two processes are intrinsically linked to mediate faithful ductular repair.

## Acknowledgements

We would like to thank the Institute of Genetics and Cancer Advanced Imaging Resource for their support with imaging and the IGC Cytometry and Single Cell Core facility for support with flow cytometry analysis and cell sorting. We would also like to thank staff at the University of Edinburgh Bioresearch & Veterinary Services for husbandry support and IGC HTPU microarray services facility for support with Reverse Phase Protein Array analysis. Fibrogen generously provided the FG-3019 and control antibodies.

## Author Contributions

AW planned and performed experiments, analysed data and edited the manuscript. PO, EC, AG, EB, LC, DW, WM, KO AB, PG, EJJ, YY produced and analysed data and generated figures for the manuscript. AML, AEPL and KD provided experimental support and technical input. TK provided support with human tissue and NC, AUB provided reagents and intellectual direction. LvdL, MMAV and MF provided intellectual input into the project. SW planned experiments, wrote and edited the manuscript and provided project level support and advice. LB led the project, funded the project, designed experiments, analysed data, and wrote and edited the manuscript.

## Data and materials availability

All data is available in the manuscript or the supplementary materials. Previously published single cell RNAseq data from this study is available from: GSE179601 and GSE243981. Previously published bulk RNA-sequencing is available at: GSE156894. Data generated in this study (Figure 2 and 3) is available at Dryad: DOI: 10.5061/dryad.qrfj6q5t1. All materials generated as part of this study will be made available upon request to the corresponding authors.

## Materials and Methods

### Human tissue

Anonymized tissue was supplied after approval by the Lothian NRS BioResource (REC reference – 20/ES/0061, approval reference – SR2185 26^th^ June 2024).

### Reanalysis of single cell RNA data from *Roos et al. 2022*

Cell count matrixes were downloaded from the GEO repository (accession number: GSE179601) and read into R (v 4.2.3) and analysed with Seurat (v 4.4.0). Cells with UMI counts < 1000 & >25,000, and with mitochondrial RNA content > 50% were filtered out from the data. Data was normalized and the top 2000 highly variable features were identified using the NormiliseData and FindVariableFeatures functions. Data was scaled with Scale data and both total number of UMIs and percentage of mitochondrial gene expression were regressed out. Cells from different donors were integrated using the FindIntegrationAnchors and IntegrateData functions to minimise confounding biological effects. Dimensionality reduction was performed using the RunPCA function, and the top 8 principal components were selected for further analysis. Hierarchal clustering was preformed using the FindNeighbors and FindClusters functions, a clustering resolution of 0.5 was used. Uniform Manifold Approximation and Projection (UMAP) plots were generated with the function RunUMAP. Cell cycle scoring was performed using the CellCycleScoring function. Differentially expressed genes from each cluster were identified using a Wilcoxon Rank Sum test with the FindAllMarkers function. Differentially expressed genes unique to cluster 2 relative to all other BRCO clusters were also found using a Wilcoxon Rank Sum test with the FindMarkers function. For both tests, only genes expressed in >20% of cells in each cluster and with a log fold change >0.25 were tested. Genes with an adjusted p value < 0.01 and an absolute fold change >0.5 were considered significantly differentially expressed. Pathway analysis of significantly differentially expressed genes was preformed using the Enrichr web tool 414.

### Reanalysis of single cell data from *Andrews et al 2024*

Raw cell count matrixes of 5’ sequenced single-cell samples were downloaded from the GEO repository (accession numbers: GSE243977 and GSE247128) and read into R (v4.2.1) and analysed with Seurat (v5.1.0). Cells were filtered to ensure mitochondrial RNA content < 25% and feature counts >1000 & <4000. Data was normalized and the top 2000 highly variable features were identified using the NormiliseData and FindVariableFeatures functions and scaled using ScaleData. Individual samples were then integrated into a single dataset using RPCA based integration through the IntegrateLayers function. Heirarchical clustering and UMAP projection was then performed as described for the Roos et al dataset above. Cell type annotations were performed in Singler (v2.10.0)^24^ using the Human Primary Cell Atlas as a reference dataset. Ligand-receptor analysis was performed in CellChat (v1.6.0)^25^. Gene signature scores for GFOGER-binding integrins were calculated using the AddModuleScore function.

### Induction and modulation of ductular regeneration in mouse

Adult mice from either the CD-1 (ECM characterisation) or C57Bl/6 (all other experiments) were used to initiate ductular regeneration. Animals were maintained in SPF environment and studies carried out in accordance with the guidance issued by the Medical Research Council in “Responsibility in the Use of Animals in Medical Research” (July 1993) and licensed by the Home Office under the Animals (Scientific Procedures) Act 1986. Experiments were performed under project license number PFD31D3D4 in facilities at the University of Edinburgh (PEL 60/6025).

All DDC injury was induced for a maximum of fourteen days, by feeding exclusively with 0.1% DDC (Sigma-Aldrich) incorporated into standard RM1 feed. For lineage tracing experiments, Keratin-19-Cre^ERT^ animals (Jax: 026925) were crossed with tdTomato (Jax: 007914) and given 4mg of Tamoxifen (Sigma-Aldrich) three times per week in 5% molecular grade ethanol and corn oil by oral gavage (throughout this paper, called *Krt19*CreER^T^; R26^LSL-tdTomato^). For knockout of *Itga2*, animals with floxed *Itga2* exon 1 were crossed into Krt19-Cre^ERT^; R26^LSL-tdTomato^ mice to generate a compound strain, *Krt19*CreER^T^; R26^LSL-tdTomato^; *Itga2*^flox/flox^.

To inhibit CTGF signalling, animals were dosed with 10mg/ml of FG-3019 (Fibrogen inc) or hIgG isotype control four times during the 14-day DDC time course. Pictilisib (MedChem Express) was reconstituted 0.5% Tween® 80 and 0.5% methylcellulose, and administered daily by oral gavage at a dose of 50 mg/kg. NXP900, provided by Neil Carragher and Asier Unciti-Broceta, University of Edinburgh was reconstituted in citrate buffer and administered at a dose of 80 mg/kg daily by oral gavage.

### Digestion and isolation of bile ducts

To isolate bile ducts from both uninjured and injured livers, dissected liver was chopped into 5-mm3 pieces and digested in DMEM/F-12 media containing Collagenase-IV (Roche) and DNASe-I (Roche). Following digestion and dissociation, bile ducts become obvious as parenchyma is digested away. Bile ducts are strained through a 70-µm filter and extensively washed in PBS to remove any residual cells. Bile ducts are then used for downstream applications.

### FACS staining and isolation

Experimental mice undergoing ductular regeneration were perfused with saline and livers were digested with collagenase and dispase to enrich for the biliary tree. Enriched bile ducts were dissociated into single cells using trypsin and stained using the antibodies detailed in supplementary table 5. Live cells (which are negative for DAPI) were identified and CD31-/CD45-EpCAM+/tdTomato+ cells were isolated for downstream applications.

### RNA isolation, cDNA synthesis and qRT-PCR

RNA was extracted from FACS isolated cells or isolated bile ducts and extracted by TRIzol RNA Isolation Reagent (Invitrogen) lysis. RNA was precipitated with chloroform and cleaned up using the RNeasy Mini Kit (QIAGEN) as per the manufacturer’s instructions. For downstream sequencing applications RNA quality (RIN score) was quantified using the Agilent 2100 Bioanalyzer with an RNA 6000 chip. A minimum RIN threshold of 8 was used for RNA-seq. Quantiect® reverse transcriptase kit (QIAGEN), including gDNA Wipeout buffer for elimination of genomic DNA, was used as per manufacturer’s instructions for cDNA synthesis. Quantitative PCR (qPCR) was performed with SYBR™ Green PCR Master Mix (Applied Biosystems™), and run in an Roche LightCycler® 480 II.

### Bulk RNA sequencing

Total-RNA samples were fragmented to a size appropriate for sequencing on an Illumina platform and first-strand cDNA was generated using the SMARTer® Stranded Total RNA-Seq Kit v2 – Pico Input Mammalian kit (Clontech Laboratories, Inc. #634411). Illumina-compatible adapters and indexes were then added via 5 cycles of PCR. Depletion of ribosomal cDNA (cDNA fragments originating from highly abundant rRNA molecules) was performed using ZapR v2 and R-probes v2 specific to mammalian ribosomal RNA and human mitochondrial rRNA. Sequencing was performed on the NextSeq 2000 platform (Illumina Inc, #20038897) using NextSeq 2000 P2 Reagents (200 Cycles) (#20046812). Libraries were combined in a single equimolar pool of nine based on Qubit and Bioanalyser assay results and run on a single P2 flow cell. PhiX Control v3 (Illumina, #FC-110-3001) was spiked in at 1% library concentration to facilitate troubleshooting in the event of any run issues.

### RNA sequencing data processing and analysis

The primary RNA-Seq processing, quality control to transcript-level quantitation, was carried out using nf-core/rnaseq v1.4.3dev (https://github.com/ameynert/rnaseq)(Thorvaldsdóttir et al., 2013). Reads were mapped to the mouse FVB_NJ_v1 decoy-aware transcriptome using the salmon aligner (1.1.0). RNA-Seq analysis was performed in R (4.0.2), Reads were summarized to gene-level and differential expression analysis was performed using the bioconductor packages tximport (1.16.1) and DESeq2 (1.28.1). A pre-filtering was applied to keep only genes that have at least 10 reads in a group and 15 reads in total. The Wald test was used for hypothesis testing for pairwise group analysis. A shrunken log2 fold changes (LFC) was also computed for each comparison using the adaptive shrinkage estimator from the ’ashr’ package.

### Immunohistochemistry and Immunofluorescence

Dissected tissues were fixed overnight in formalin at 4°C, embedded in paraffin and were sectioned at 4µm. Following antigen retrieval (see Supplementary materials table 5), tissue sections were incubated with antibodies as detailed in Supplementary materials table 5. Fluorescently stained tissues were counterstained with DAPI prior to imaging. Colorimetric stains were counterstained with Haematoxylin and mounted with DPX.

### FunGI clearing and confocal imaging

Regenerating, DDC-treated tissues were sectioned at 200 µm using a Krumdieck Tissue Slicer and fixed for 45 min in formalin and then cleared using FUnGI clearing as previously described (Dawson et al., 2020; Younger et al., 2022). Histological tissues were scanned using a Nanozoomer, using a Nikon A1R or Leica Stellaris confocal microscope and were analysed using either FUJI, Imaris or QuPath.

### Cross correlation of AFM with confocal imaging

**A**tomic force microscopy (AFM) measurements were performed at the Centre for Science at Extreme Conditions, University of Edinburgh. Briefly, 20μm cryosections were rehydrated and 100μm areas around portal veins were scanned on an JPK NanoWizard 4XP BioScience atomic force microscope. Following AFM measurements cryosections were fixed in 10% neutral buffered formalin and incubated overnight at 4°C with primary antibody (supplementary table 5), followed by incubation with AlexaFluor-conjugated secondary antibodies at room temperature. During imaging, the correct area of AFM measurement was identified on brightfield imaging and 10μm confocal z-stack images were taken and a brightfield image included for image registration.

Image registration was performed using multimodal image registration on NIS Elements software, on brightfield images taken from both the AFM and confocal microscopes; ten fiducial marks were made for each slide using the portal vein area and porphyrin plugs, and images from AFM and immunofluorescence were merged to a confocal stack. All further analysis was performed in QuPath using or ImageJ (JACoP Plugin^38^).

Histological analysis of collagen staining was performed using Qupath. Utilising a pixel classification script to determine H scores based on DAB intensity. To quantify staining intensity based on pixel classification, the H-score (which is scaled between 0-300) is calculated using the following formula:

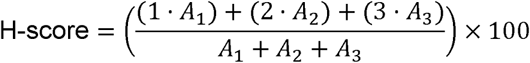

where:

- *A*_1_ is the area classified as low positive in µm^2^,
- *A*_2_ is the area classified as moderate positive.
- *A*_3_ is the area classified as strong positive.

In our analysis banding of *A^1^-A^3^* was performed by splitting the histogram of pixel wise intensity into three, this gave us the following values. *A^1^ =* 0.25*, A^2^ =* 0.4*, A^3^ =* 0.6.

### Measuring bile duct migration *in vivo*

To quantify the migratory distance of bile ducts from the portal vein, portal veins with an area of between 1000 and 5000 μm2 were included when quantifying bile duct migration in DR. The liver area surrounding portal veins was mapped in 5μm steps up to 100μm expansion from the original portal vein. The biliary content of each concentric ring, as measured by EpCAM-positive staining, was calculated.

### Culture of human BEC cell line H69

H69s were cultured Advanced DMEM/F12 (Gibco), 1% GlutaMAX™ (Gibco), 1% penicillin/streptomycin, 5% heat-inactivated sterile-filtered foetal bovine serum (Life Science Group), 1% lipid mixture 1, chemically defined (Sigma-Aldrich),1x MEM vitamin solution (Sigma-Aldrich), 1x insulin-transferrin-selenium (Gibco), soybean trypsin inhibitor 50 μg/mL (ATCC), 3, 3’ 5-triiodo-L-thyronine (T3)(Sigma-Aldrich), bovine pituitary extract 13.4 μg/mL (1x)(Life Technologies), dexamethasone 0.393 μg/mL (1x)(Sigma-Alrich), epidermal growth factor (EGF) 25 ng/mL (1x, stock made up in PBS with 1X BSA)(Sigma-Aldrich), forskolin 4 μg/mL (Ascent Scientific). Ten drops of NaOH 3,4N were added to every 500mL media, which was then filtered through a 0.22 μm membrane.

### siRNA of integrins *in vitro*

RNA interference was carried out using Lipofectamine® RNAiMAX Reagent (Invitrogen™) and Opti-MEM™ (Gibco™). All RNAi (against Itga2 and Itgb1, Supplementary table 4) experiments were carried out as per manufacturer’s instructions RNAi was carried out 72 hours prior to the whole protein lysate extraction or 48 hours prior to the start of experiments, at which point the media including siRNA was refreshed.

### Culture and differentiation of BRCOs

BRCO cultures were generated as previously published. Briefly, ICOs were generated from cryopreserved patient biopsies from healthy livers and cultured for three days in BME domes with SEM. Once successfully established, SEM was replaced with EM to support normal organoid expansion. Organoids were maintained in EM but did not exceed passage nine.

When ICOs reached approximately 80% confluency within the BME dome, they were split at a 1:10 ratio into fresh domes to ensure ∼20% confluency. For BRCO generation, wells were seeded, and EM was added to each well. After three days of culture in EM, ICOs were present, and the media was switched to BM. Each well received 1 mL of BM, containing either 50 µM TCI-15 or a DMSO control. Brightfield images of organoids were taken before BM addition and every 24 hours thereafter. BM containing TCI-15 or DMSO was refreshed every 2–3 days. To assess branching, the perimeters of individual organoids were measured and defined as the deviation from a perfect circle, which would measure 1.

### Protein isolation and Immunoblotting

All whole protein lysate was obtained by dissociating tissue in 1x RIPA lysis and extraction buffer (Millipore), supplemented with 1x cOmplete™, Mini, EDTA-free Protease Inhibitor Cocktail (Roche) and 1x PhosSTOP™ (Roche). Protein lysates (5-20μg) were mixed with LDS sample buffer and sample reducing agent (Invitrogen) and denatured at 70°C. Protein samples were separated using Bis-Tris, 4–12% (Invitrogen) with MOPS SDS running buffer (Invitrogen) at 150V. Wet transfer on either nitrocellulose or methanol-activated PVDF 0.45 μm transfer membrane were performed using transfer buffer (Invitrogen with 20% methanol). Membranes were blocked with 3% bovine serum albumin (BSA) (Sigma-Aldrich) in Tris-buffered saline, 0.1% Tween® 20 detergent (TBS-T).

Immunoblots were probed with primary antibodies as detailed in supplementary table 6 and HRP-conjugated secondary antibodies in blocking buffer. Target proteins were detected with Pierce™ ECL Western blotting substrate (Thermo Scientific) and imaged on an Amersham™ ImageQuant™ 800 (Cytiva). Western blot data was quantified in ImageJ (Fiji).

### Quantifying proliferation and migration *in vitro*

Proliferation and migration assays were performed on the Incucyte® Live-Cell Analysis instrument (SARTORIUS). All experiments were carried out in 96-well plates, coated with collagen I unless stated otherwise. For proliferation assays, cells were seeded at between 1000-5000 cells/well. Migration assays were performed on confluent wells, seeded at 10,000 cells per well. Migration was assessed by wound healing, using a Woundmaker as per manufacturer’s instructions. All analyses were performed using Incucyte 2022B Rev2.

**Supplementary Table 4:**
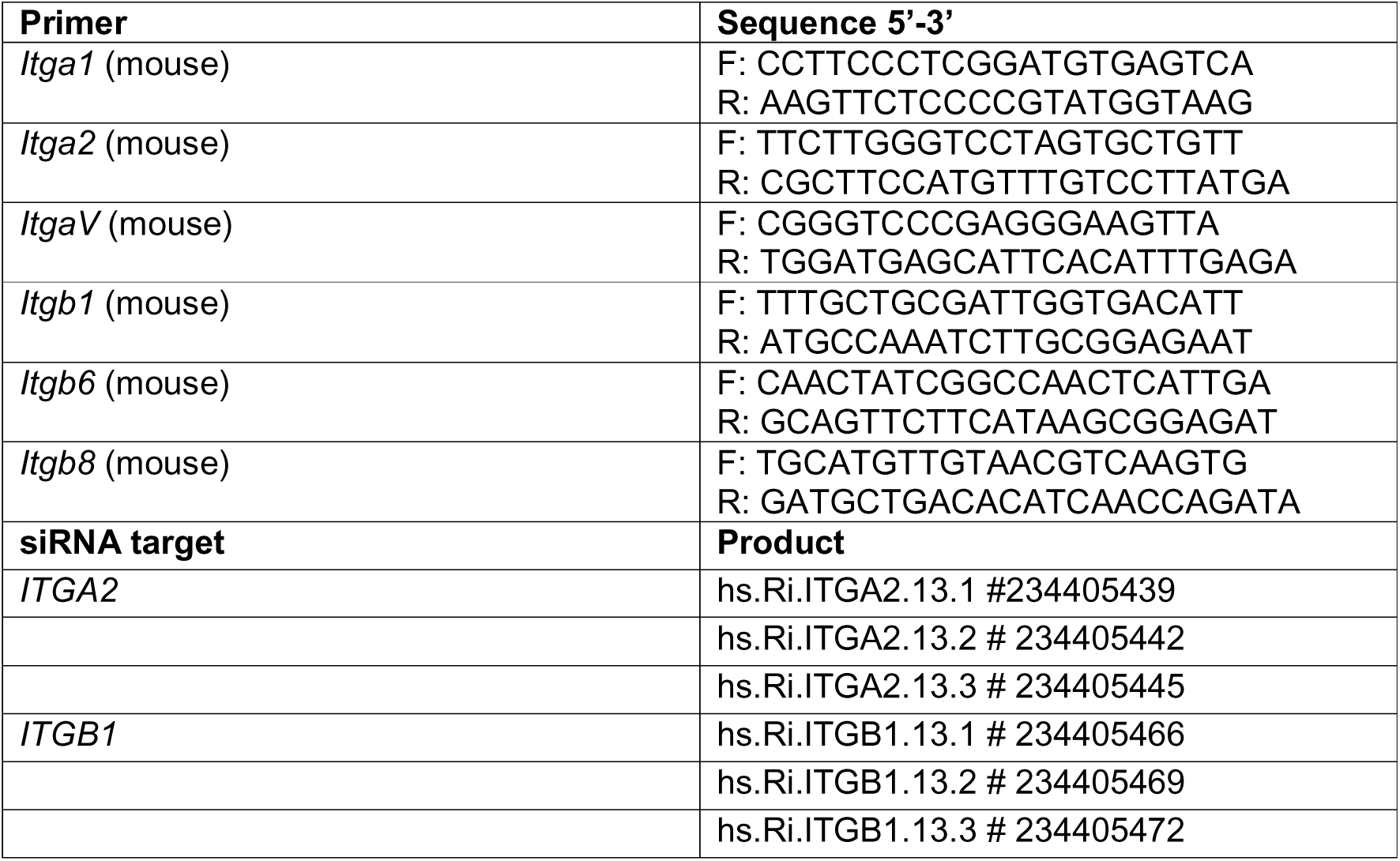
qRT-PCR primers in this study:

**Supplementary Table 5:** Immunohistochemistry antibodies used in this study.

| Immunohistochemistry: |  |  |
| --- | --- | --- |
| Primary Antibodies: |  |  |
| Target | Antigen retrieval | Antibody and clone |
| Integrin α1 | TE pH9, 4min pressure cooker | Abcam AB243032 [CL7207] |
| Integrin α2 | Sodium citrate, 10min microwave | Abcam AB133557 [EPR5788] |
| Integrin β1 | Sodium citrate, 10min microwave | Abcam AB179471 [EPR16895] |
| Collagen I | Sodium citrate, 10min microwave | Southern Biotech 1310-01 |
| Collagen III | Sodium citrate, 10min microwave | Abcam AB7778 |
| Collagen IV | Sodium citrate, 10min microwave | Southern Biotech 1340-01 |
| Laminin | Sodium citrate, 10min microwave | Abcam AB11575 |
| Fibronectin | Sodium citrate, 10min microwave | Invitrogen MA5-11981 |
| Keratin 19 | TE pH9, 4min pressure cooker | DHSB Troma-III-c |
| EpCAM | TE pH9, 4min pressure cooker | Abcam AB71916 |
| Total FAK | TE pH9, 4min pressure cooker | Cell Signalling Technologies #3285 |
| pFAK Y397 | TE pH9, 4min pressure cooker | Invitrogen 44-624G |
| pPYK2 | TE pH9, 4min pressure cooker | Cell Signalling Technologies #3291 [Y4B2] |
| A-SMA | TE pH9, 4min pressure cooker | Sigma-Aldrich A2547 [1A4] |
| F4/80 | Proteinase K in TE, 10min RT | Abcam AB6640 [CI-A3-1] |
| CD45 | TE pH9, 4min pressure cooker | Novus NBP2-34528 |
| Akt | TE pH9, 4min pressure cooker | Cell Signalling Technologies #4691 |
| pAkt S473 | Citrate buffer, 20min microwave | Cell Signalling Technologies |
|  |  | #4060 [D9E] |
| tdTomato | TE pH9, 4min pressure cooker | SICGEN mCherry AB0081-200 |
| PCNA | TE pH9, 4min pressure cooker | Abcam AB29 [PC10] |
| *Secondary Antibodies:* |  |  |
| Anti-goat | 488 | Thermo Scientific A-11055 |
|  | 594 | Thermo Scientific A-11058 |
| Anti-mouse | 488 | Thermo Scientific A-11029 |
|  | 594 | Thermo Scientific A-21203 |
|  | 647 | Thermo Scientific A-31571 |
| Anti-rabbit | 488 | Thermo Scientific A-11034 |
|  | 594 | Thermo Scientific A-21207 |
|  | 647 | Thermo Scientific A-31573 |
| Anti-rat | 488 | Thermo Scientific A-21208 |
|  | 594 | Thermo Scientific A-21209 |

**Supplementary Table 6:** Western blotting and FACS antibodies used in this study.

| **Western Blotting** |  |
| --- | --- |
| *Primary Antibodies* |  |
| **Target** | **Antibody** |
| Total FAK | Cell Signalling Technologies #3285 |
| pFAK Y397 | Cell Signalling Technologies #8556 [D201B] |
| pFAK Y576/7 | Cell Signalling Technologies #3281 |
| pFAK Y925 | Cell Signalling Technologies #3284 |
| Total Src | Cell Signalling Technologies #2110 [L4A1] |
| pSrc Y416 | Cell Signalling Technologies #59548 [E6G4R] |
| Akt | Cell Signalling Technologies #4691 |
| pAkt S473 | Cell Signalling Technologies #4060 [D9E] |
| B-tubulin | Cell Signalling Technologies #2128 [9F3] |
| GAPDH | Cell Signalling Technologies #5174 [D16H11] |
| *Secondary Antibodies* |  |
| anti-goat HRP | Vector Laboratories BA-5000 |
| anti-mouse HRP | Vector Laboratories PI-2000 |
| anti-rabbit HRP | Vector Laboratories PI-1000 |
| anti-rat HRP | Abcam ab97090 |
| **Flow cytometry** | **Antibody.** |
| **Target** |  |
| CD31 | CD31 PE-Cy7 (BD Pharmingen #561410) |
| CD45 | CD45 PE-Cy7 (BD Pharmingen #552848) |
| CD326 (EpCAM) | CD326 (Ep-CAM) APC (BioLegend® #118214) |
| CD49b (ITGA2) | α2-integrin-APC-Cy7 SouthernBiotech #1806-19 |

## List of Supplementary Tables and Figures

Supplementary Table 1: Ligand-Receptor interaction outcomes between fibroblasts and BECs in human biliary disease.

Supplementary Table 2: Details of human PSC samples used in this study.

Supplementary Table 3: Differentially expressed genes in ITGA2 high versus low BECs.

Supplementary Table 4: Differentially expressed genes in BRCO cluster 2.

Supplementary Table 5: Differentially expressed genes following *Itga2*-deletion.

Supplementary Table 6: qRT-PCR primers in this study:

Supplementary Table 7: Immunohistochemistry antibodies used in this study.

Supplementary Table 8: Western blotting and FACS antibodies used in this study.

**Supplementary Figure 1:**
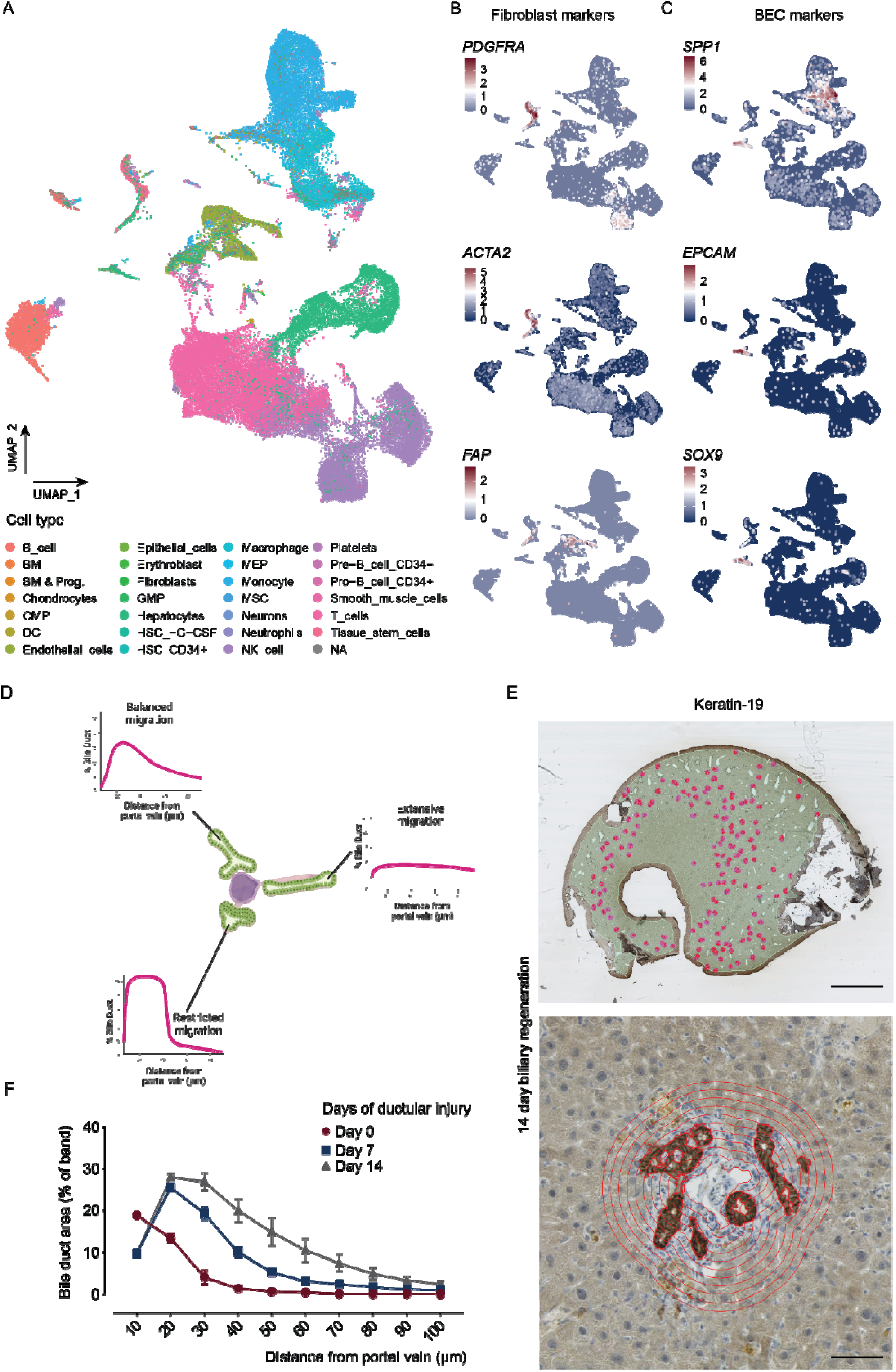
The formation of a collagen niche is a hallmark of ductular regeneration. **A.** SingleR analysis and UMAP projections of single cell data from *Andrews et al., 2024* demonstrating a wide range of cell types from normal human liver and patients with Primary Sclerosing Cholangitis and Primary Biliary Cholangitis. **B.** Cell subtype validation of Fibroblast markers (*PDGFRA*, *ACTA2* and *FAP*) and **C.** BEC markers (*SPP1*, *EPCAM*, *SOX9*). **D.** A schematic representation of how ductular regeneration can be modulated. In normal, physiological regeneration, bile ducts migrate away from the portal tract. Experimental perturbation could lead to extensive or restricted migration. **E.** Representative images of the distribution of Keratin-19-positive portal tracts analysed (upper panel, scale bar = 0.5cm) and the automated radial quantification, using the portal vein as an anchor point, to measure distance travelled (lower panel, scale bar = 200μm). **F.** Measurements of the distance of bile ducts from portal veins (N=3 biological replicates per time point) in normal liver (Day 0, red line) or following either 7 or 10 days of ductular regeneration (blue and grey lines, respectively).

**Supplementary Figure 2:**
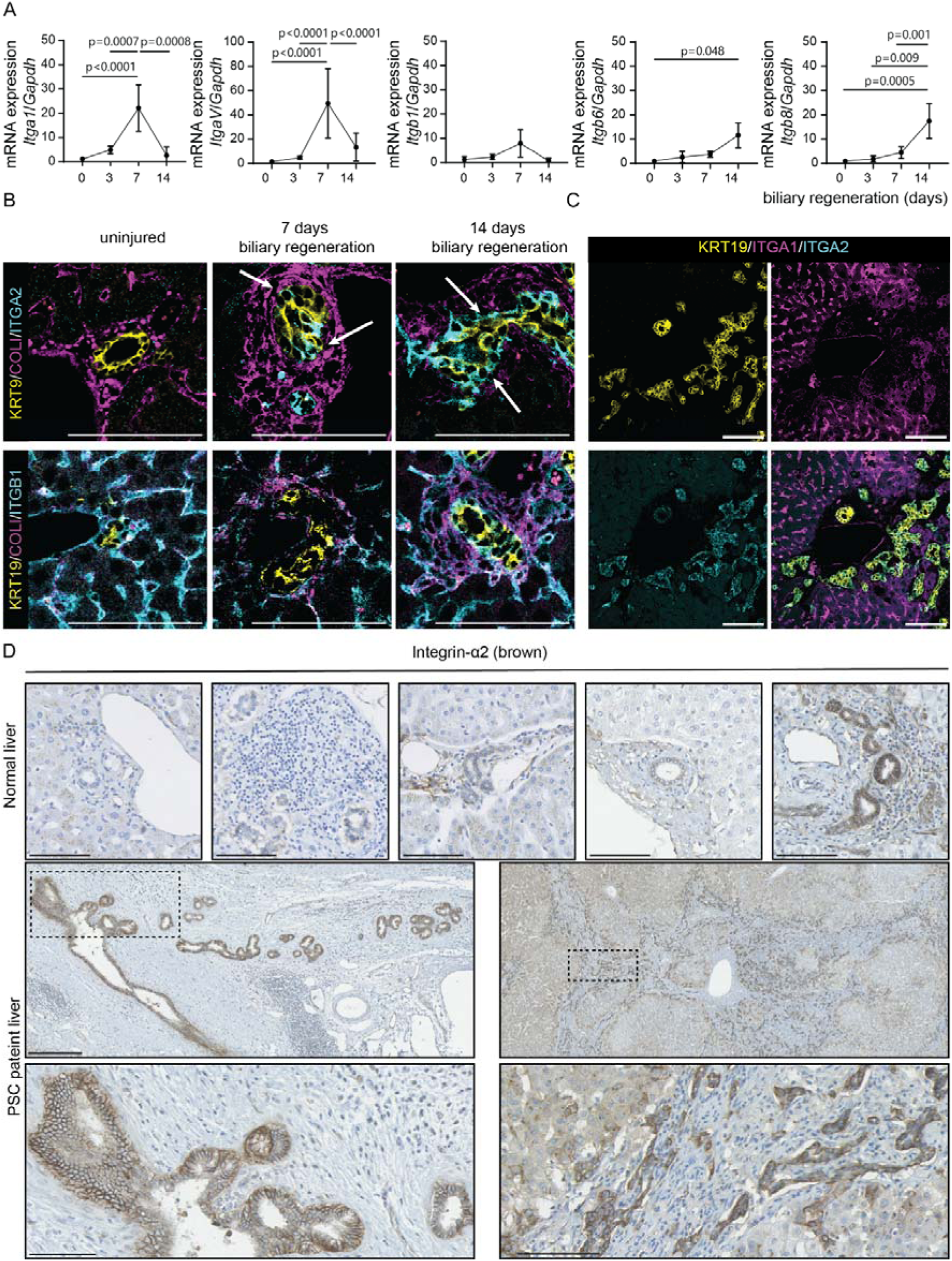
Specific integrins are expressed during ductular regeneration: **A.** mRNA expression of *Itga1*, *ItgaV*, *Itgb1*, *Itgb6* and *Itgb8* in isolated bile ducts from mice undergoing ductular regeneration (N=4 animals per time point). **B.** Immunofluorescent staining of Keratin-19 positive bile ducts (yellow), collagen-I (magenta) and integrin-α2, *upper panels* or integrin-β1, *lower panels*, cyan. Scale bar = 200μm. **C.** Immunofluorescence of mouse livers following 14 days of ductular regeneration stained for Keratin-19 positive bile ducts (yellow), integrin-α1 (magenta) and integrin-α2, cyan. Scale bar = 150μm. **D**. Immunohistochemistry of integrin-α2 in normal human tissue and in PSC-patient tissue. Scale bar = 250μm upper panels and lower panel, scale bar = 100μm.

**Supplementary Figure 3:**
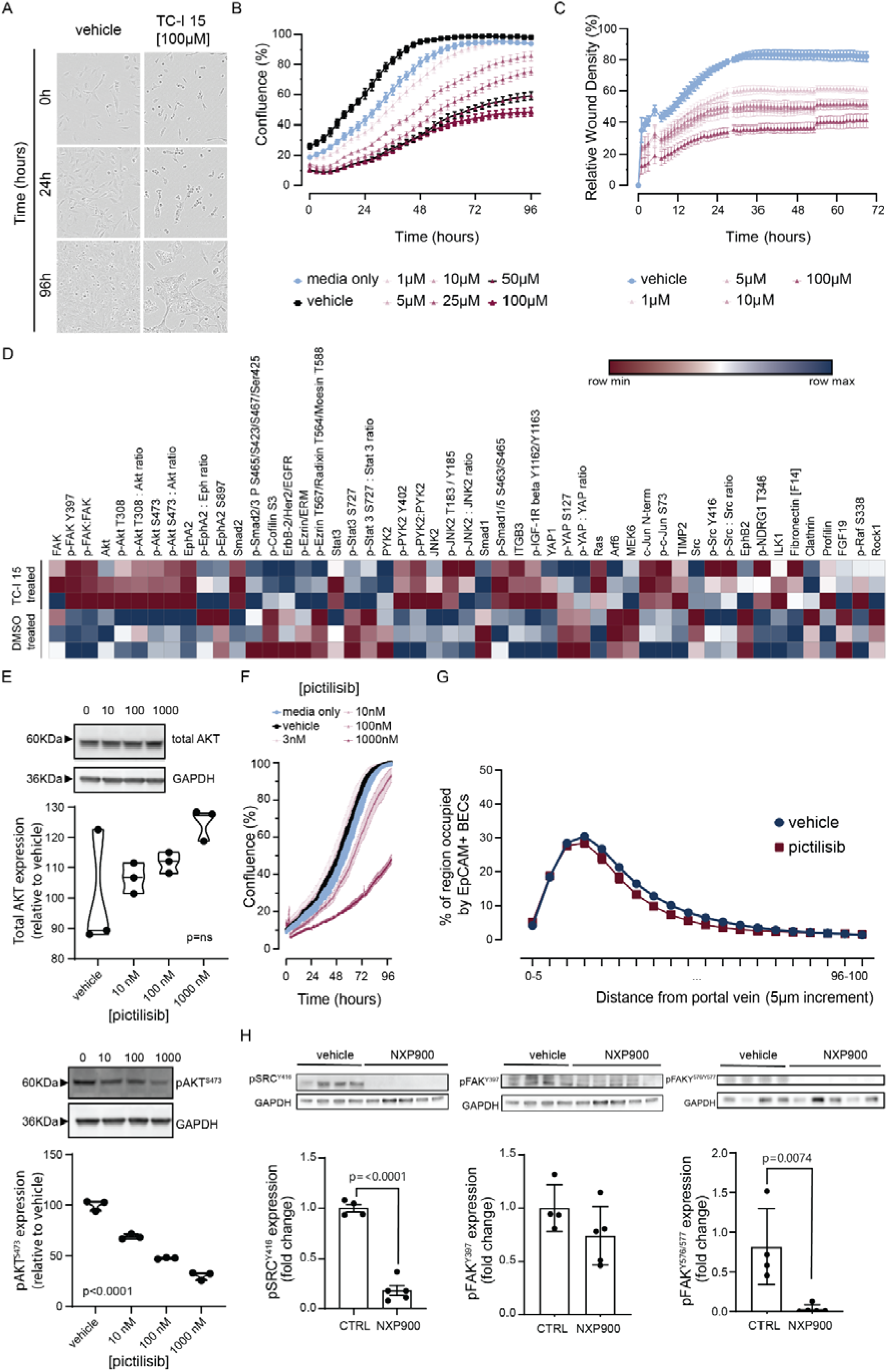
Identifying integrin-dependant signals that regulate ductular regeneration. **A.** Representative timelapse photomicrographs of hBECs (H69 cells) treated with vehicle or 100μM integrin-α2β1 inhibitor, TC-I 15 over 96h. **B.** cell growth and **C.** wound-closure of hBECs following treatment with varying doses of TC-I 15 (n=3 technical replicates per condition). **D.** Reverse Phase Protein Array in hBECs following treatment with TC-I 15 (N=3 technical replicates per group). E. Immunoblots (upper panels) for total AKT and phosphorylated AKT^S473^ in hBECs treated with increasing concentrations of the Pi3K-inhibitor, Pictilisib. (N=3 technical replicates). **F.** Timelapse imaging of cellular growth of hBECs following treatment with Pictilisib at different concentrations. (N=3 technical replicates per concentration). **G.** Quantification of bile duct migration following 14 days of biliary regeneration in mice treated with Pictilisib or vehicle alone (N=4 mice per group). **H.** Immunoblot and quantification of SRC^Y416^, FAK^Y397^, FAK^Y576/577^ and GAPDH in animals with ductular regeneration treated with vehicle or NXP900 (N=4-5 mice per group).

**Supplementary Figure 4:**
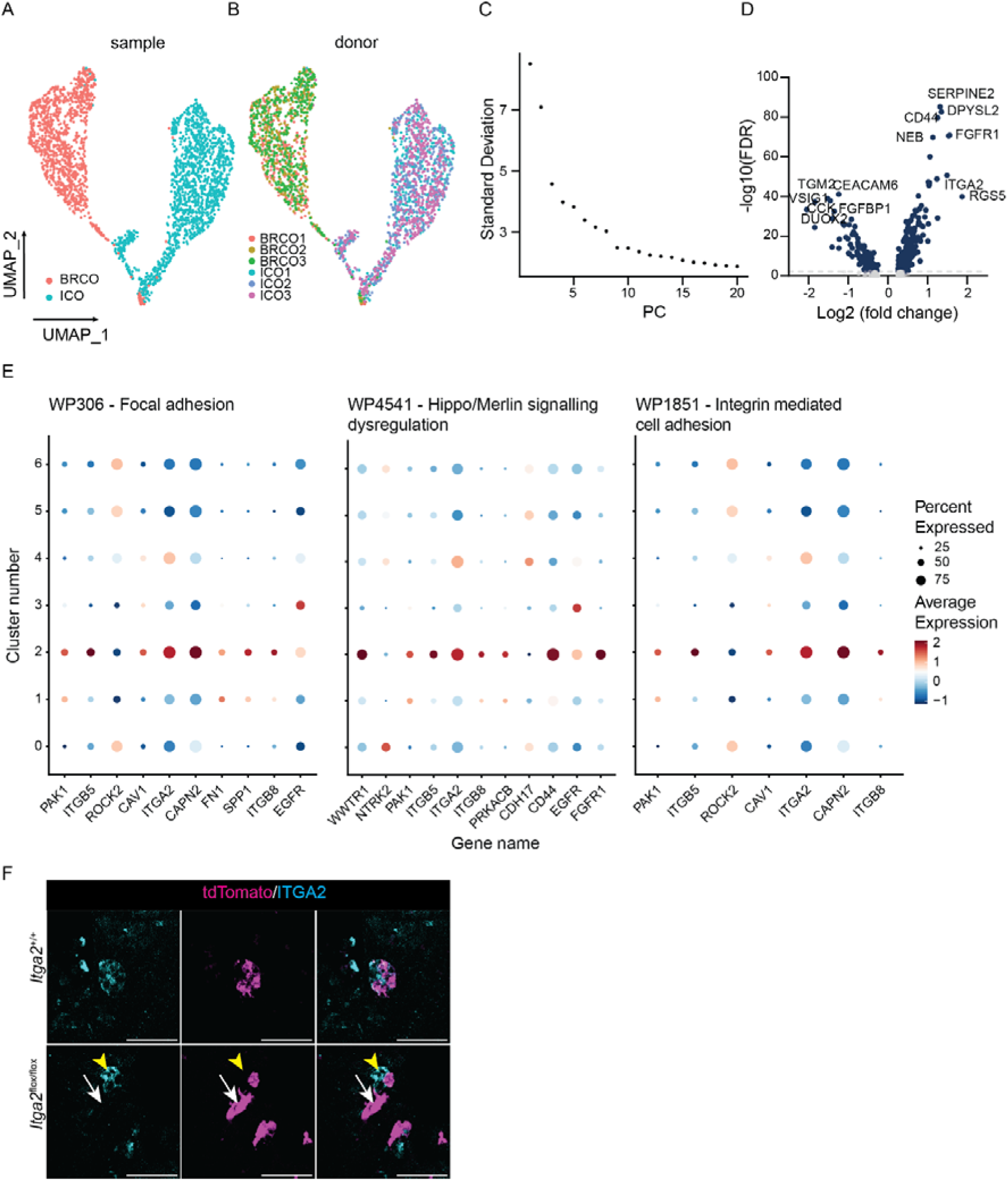
Modulating integrin limits ductular morphogenesis. **A.** UMAP projections of data generated by *Roos et al., 2022* demonstrating that intrahepatic cholangiocyte organoids (ICOs, teal) cluster differently to differentiated branching cholangiocyte organoids (BRCOs, salmon). **B.** Donor sample projected onto the UMAP demonstrating consistency in cluserting of ICO and BRCO experimental groups. **C.** Elbow plot used to define the number of Seurat Clusters in single cell data. **D.** DEGs enriched in cluster 2 relative to all other clusters expressed as a volcano plot. **E.** Cluster-by-cluster mRNA expression from single cell data using GOTerm-specific gene sets for WP306 – Focal adhesion, WP4541 – Hippo/Merlin Signalling dysregulation and WP1851 – integrin mediated cell adhesion. **F.** Immunohistochemistry for tdTomato (magenta) and ITGA2 (cyan) in *Krt19*-Cre^ERT^; *Itga2*^+/+^ or *Itga2*^flox/flox^ transgenic mice. Yellow arrow demarcates ITGA2 staining and white arrow, tdTomato positive cells in which ITGA2 has been lost. Scale bar = 100μm.

